# Observation and simulation of chemically mediated searches in marine zooplankton

**DOI:** 10.1101/2025.07.30.666938

**Authors:** Spencer J. Franks, Clayton Vondriska, Peter Hinow, Paul C. Sikkel

**Author notes:** These authors contributed equally to this work (Spencer J. Franks, Clayton Vondriska).

## Abstract

Chemical communication is one of the most important capabilities of zooplankton, yet it remains understudied. We report observations of marine parasitic isopods *Gnathia marleyi* and coral larvae *Orbicella faveolata* in an aquatic olfactometer where individuals can respond to chemical signals from host fish species and crustose coralline algae, respectively. We complement these observations with a gradient search model inspired by the theory of *Braitenberg vehicles*. The mathematical model allows the agent to switch between periods of motion and periods of rest, guided by the behavior of *G. marleyi*. We further implement a temporary coupling of the agent to the signal gradient, interspersed with periods of seemingly aimless wandering. We find a favorable comparison between the experimental and simulated trajectories. This study gives us further insight on the chemosensory behavior of ecologically important marine invertebrates using novel instrumentation and methodology and is the first to attempt to model the chemosensory behavior of these two groups of aquatic invertebrates. Further research should seek to observe and model their behavior in three-dimensional space.

## Introduction

Many animals live in complex olfactory landscapes and exploit chemical signals to locate food and mates, explore habitat, and avoid predators. Chemical signals are of particular importance in aquatic ecosystems, where other stimuli such as visual or acoustic signals can be easily inhibited by environmental factors [1–3]. In aquatic systems, small invertebrates dominate biomass and biodiversity [4]. Compared to their terrestrial counterparts, the aquatic microfauna is severely understudied. This can be attributed to a combination of their small size and crypsis, the difficulties associated with humans operating in aquatic environments, and their perceived lack of direct effects on humans. In the field of sensory ecology, some of the most well studied microfauna include ectoparasites of humans and other vertebrates such as mosquitoes and ticks. These organisms directly take nutrients from their host and can act as vectors for blood-borne pathogens. In their respective ecosystems, aquatic organisms are not less significant and can also have effects on human populations, although they may not be as obvious [5]. In aquatic systems microfauna can often negatively affect important food sources of humans (e.g. salmon louse and aquaculture fish, [6]) or can be extremely influential in their ecosystems (e.g. reef building corals). Here, we study the chemosensory behavior of two aquatic invertebrates, an ectoparasitic gnathiid isopod and a larval stage of a Scleractinian coral.

Complementing these observations, we use a mathematical model for *Braitenberg vehicles* [7] following earlier work [8]. We adapt it to our purposes by introducing a gradient search mechanism and adding stochastic components modeling rest-and-run behavior, random perturbations, and a possible loss of vigilance. Originally designed by Valentino Braitenberg (Italian-German neuroscientist, 1926-2011) as thought experiments in psychology [7], Braitenberg vehicles react to sensor input and can exhibit remarkably complex behaviors. A natural inspiration is the sense of hearing in birds or mammals where the distance between the two ears allows the brain to detect the position of a source of sound [9, 10]. Insects [11] and snakes [12] can detect even small concentration gradients of a chemical with a small size detection apparatus. Similarly, gnathiids possess paired anterior antennae that are likely to be used to detect chemical stimuli, as other crustaceans [13, 14]. An exciting related field of research is mobile odor tracking robots that can detect sources of hazardous chemicals, so-called “artificial noses” [15, 16]. Similarly, Shaikh and Rañó [17] study Braitenberg vehicles as computational tools for neuroscience research. Our mathematical model is capable of replicating exploratory and active search behavior. Although our observations were made in the two-dimensional setting of the olfactometer, they are useful for studying the reaction of marine micro fauna to the presence of chemical signals. Furthermore, we simulate behaviors in time-dependent signal landscapes, paving the way for a better understanding of the chemical ecology of aquatic invertebrates.

## Study organisms

### Ectoparasitic gnathiid isopods

Gnathiid isopods occur in temperate, polar, and tropical oceans, from tide pools to deep ocean [18–20]. They share similarities with blood-feeding arthropods on land, such as ticks or mosquitoes. These small (typically 1-3 mm) animals differ from other marine external ectoparasites in two fundamental ways. First, since adults do not feed, they are only parasitic during each of the three juvenile phases (instars). Second, except for species that infest sharks and rays, they associate only temporarily with hosts, spending most of their life on the substrate. Thus, they have been referred to as “temporary external ectoparasites”, “protelean parasites”, and “micropredators”. First-stage parasitic juveniles emerge from the substrate, feed on a single host fish, and when engorged, return to the substrate and molt into their next stage. This cycle is repeated twice [18, 21]. Gnathiids, therefore, must navigate complex environments to locate a different host at each stage [22]. After the final blood meal, third-stage juveniles metamorphose into nonfeeding adults that live in the benthos, reproduce, and then die.

In this study, we use *Gnathia marleyi* as our representative gnathiid isopod. *G. marleyi* is a common gnathiid found in coral reef habitats in the eastern Caribbean [14]. It is known to parasitize more than 80 species of reef fish [23,24] and has been documented to travel up to 3 m (*>* 10^3^ body lengths) into the water column to feed on a host [25].

Like most tropical gnathiids, it is most active at night [26, 27] and has been shown to rely on chemical signals to locate fish hosts [28, 29].

### Coral larvae

Scleratinian (hard) corals occur primarily in shallow tropical ocean waters, declining in biodiversity as latitude increases and depth increases [30]. Corals are the most important biological substrate on coral reefs and are critical to providing shelter and food for reef organisms [31]. They also provide important structural complexity to reefs, helping moderate competition and predation interactions by providing refuge space to reef-associated fishes and invertebrates [32]. Although most corals are completely sessile as adults, coral larvae are free-living. Most coral species are synchronous broadcast spawners, releasing their male and female gametes in association with the lunar cycle [33]. After fertilization, coral larvae must quickly locate a suitable substrate on which to settle before metamorphosing into their adult polyp form. One of the most common settlement substrates chosen by corals is crustose coralline algae (CCA), a calcareous red algae [34, 35].

In this study, we used *Orbicella faveolata* as our representative coral species. *O. faveolata* is a slow-growing star coral widely found throughout the Caribbean [36]. Because of their wide distribution and long life spans *O. faveolata* is often considered one of the most important reef-building corals. *O. faveolata* release their gametes in September and October, with fertilization and settlement occurring shortly thereafter [33].

## Materials and Methods

All *G. marleyi* were wild caught on the reefs of St. Thomas, United States Virgin Islands (USVI) using lighted plankton traps similar to those described in [37]. Briefly, lighted plankton traps were made of a PVC tube with an inward facing funnel on one side and a plankton mesh covering the other. Within the trap is an underwater flashlight that shines light out of the funnel onto a portion of the reef. These traps were left undisturbed overnight, with light that attracts the plankton into the trap and the plankton mesh allowing water to flow within the trap without allowing the plankton to escape. The plankton within the trap was then classified and the unfed gnathiids were separated for behavioral trials. The remaining plankton was returned to the capture site. The host chosen for chemical stimuli was the Beaugregory damselfish (*Stegastes leucostictus*). Beaugregory damselfish are a common host of *G. marleyi* that are abundant throughout the USVI. Previous studies have suggested that *G. marleyi* is responsive to chemical cues originating from the host [29]. All fish were collected using modified cast nets and returned to the capture site after the experiments.

*O. faveolata* gametes were obtained from adult colonies using 50 mL Falcon tubes attached to conical mesh nets. Once about 20 % of the tube was filled with gamete bundles, the tubes were removed, capped, and brought to a boat for processing. The gametes from multiple colonies were then mixed and placed in flasks for transport. Gametes were held in the Rosenstiel School of Marine, Atmospheric, and Earth Sciences Laboratories in 27 °C filtered seawater (FSW) until larval development and movement was observed. Water changes were performed daily on the larval cultures. Coral gametes were collected under the Florida Keys National Marine Sanctuary permit FKNMS-2024-178. The chemical stimuli chosen for *O. faveolata* was CCA. Like many coral species that associate with CCA, chemical signals originating from CCA can induce settlement and improve settlement success [34, 35].

For experiments, we used an olfactometer as previously described [38, 39]. This olfactometer allows an individual organism to move freely throughout a choice arena while being exposed to different zones containing separate sources of water [39]. For these experiments, one quadrant of the choice arena contained water that had the chemical signals of an individual Beaugregory damselfish while the other three quadrants contained unconditioned seawater. The water was conditioned with the chemical signals of the damselfish by placing an individual directly into the test cue tank of the olfactometer for the duration of the experiment. The water was set to flow from each inlet of the choice arena at 10 ml/min, with each quadrant converging and flowing out a central outlet.

For each trial, the water flowed through the choice arena for at least 5 minutes, ensuring that each quadrant is properly filled with its source water. Then an individual juvenile unfed gnathiid isopod was introduced to the center of the choice arena and allowed to move freely for 10 minutes. Throughout the duration of the trial, the choice arena was covered with a light-proof cover with infrared lights illuminating the choice arena. Previous experiments have shown that *G. marleyi* is not reactive to infrared lighting in laboratory experiments [29]. All trials were recorded for their entire duration using a Basler acA2040-um NIR camera (Basler AG, Germany). Data were extracted from videos using ImageJ [40] and EthoVision XT software [41]. The videos were digitized and the positional data along with the python codes are available at doi.org/10.6084/m9.figshare.29671754.

### Differential equation models of Braitenberg vehicles

Rañó [8] proposed an ordinary differential equation (ODE) model for Type 2 and 3 Braitenberg vehicles in two dimensions. The state of the agent is given by its location (*x, y*) in space and its direction *θ* ∈ [0, 2*π*) which is the angle against the positive *x*-axis. The model is given by the three ODEs [8]

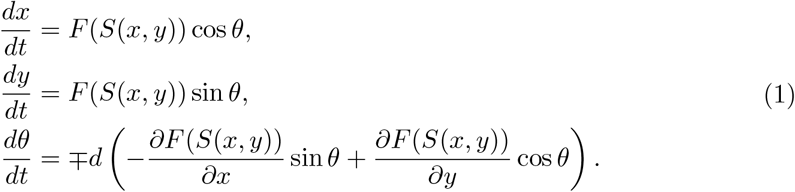

Here, *S*(*x, y*) ≥ 0 denotes the signaling landscape. For the numerical simulations in Figure 3 we use the Gaussian signal

**Figure 1.**
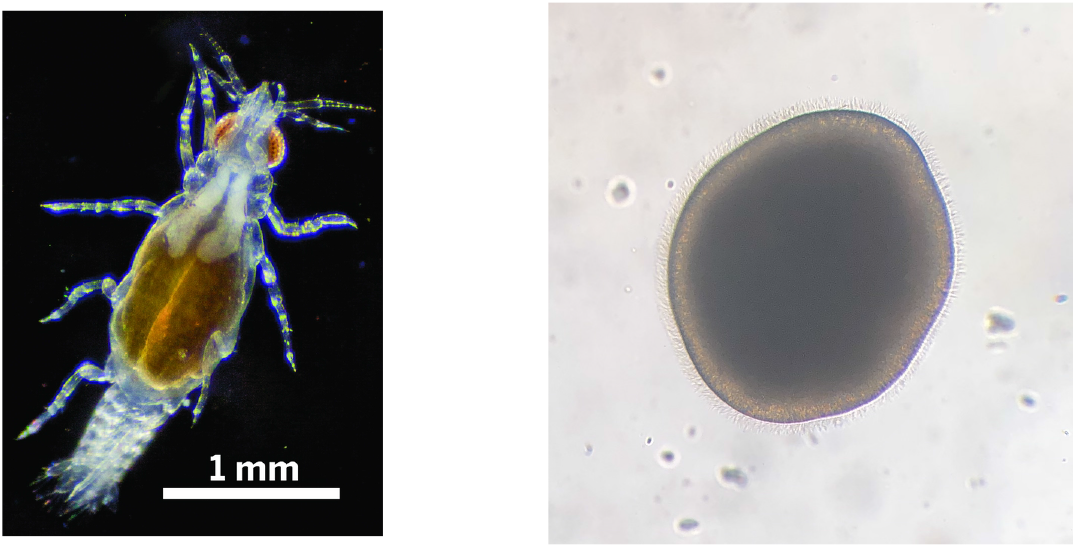
**(Left)** Image of a juvenile *Gnathia marleyi* (John Artim, Arkansas State University). **(Right)** Image of a larval *Orbicella faveolata* (Liv Liberman, Revive and Restore); the diameter is approximately 0.5 mm.

**Figure 2.**
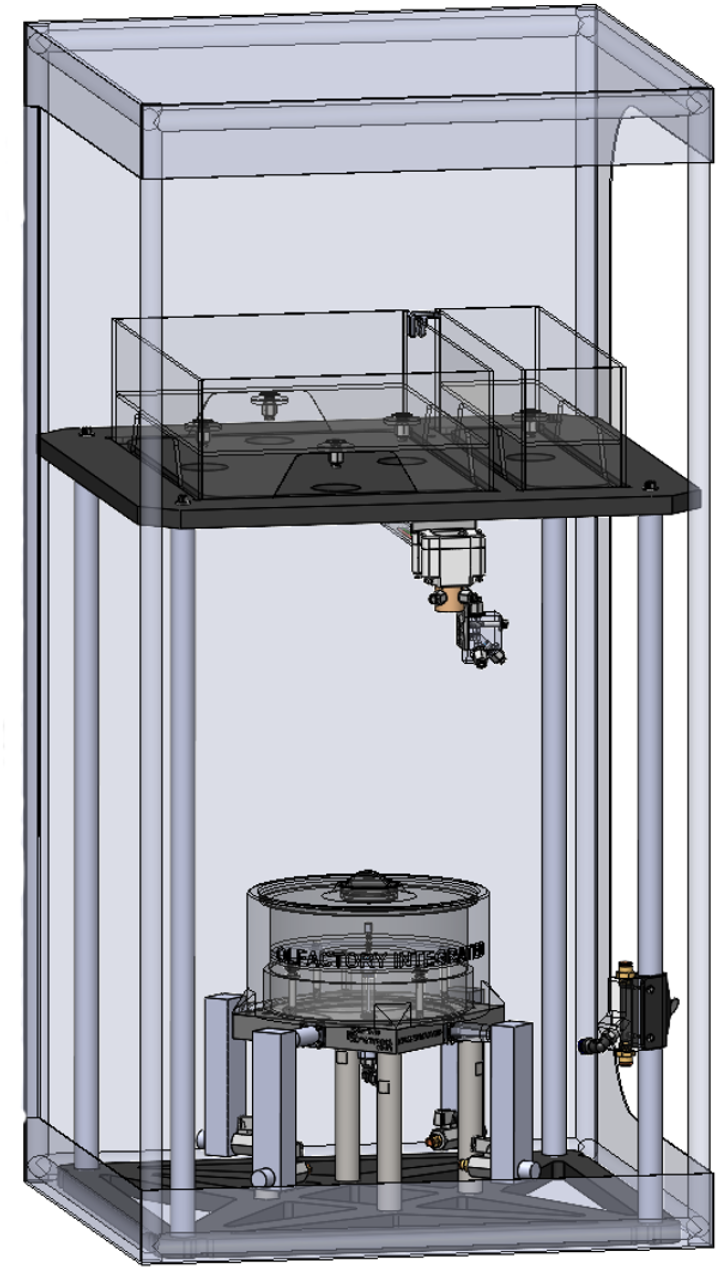
Schematic representation of the aquatic olfactometer. Water sources are located above a choice arena. The choice arena is covered by a light proof cover that is internally illuminated using infrared lighting. A high-speed infrared camera can record within the choice arena. For more details on the components see [39].

**Figure 3.**
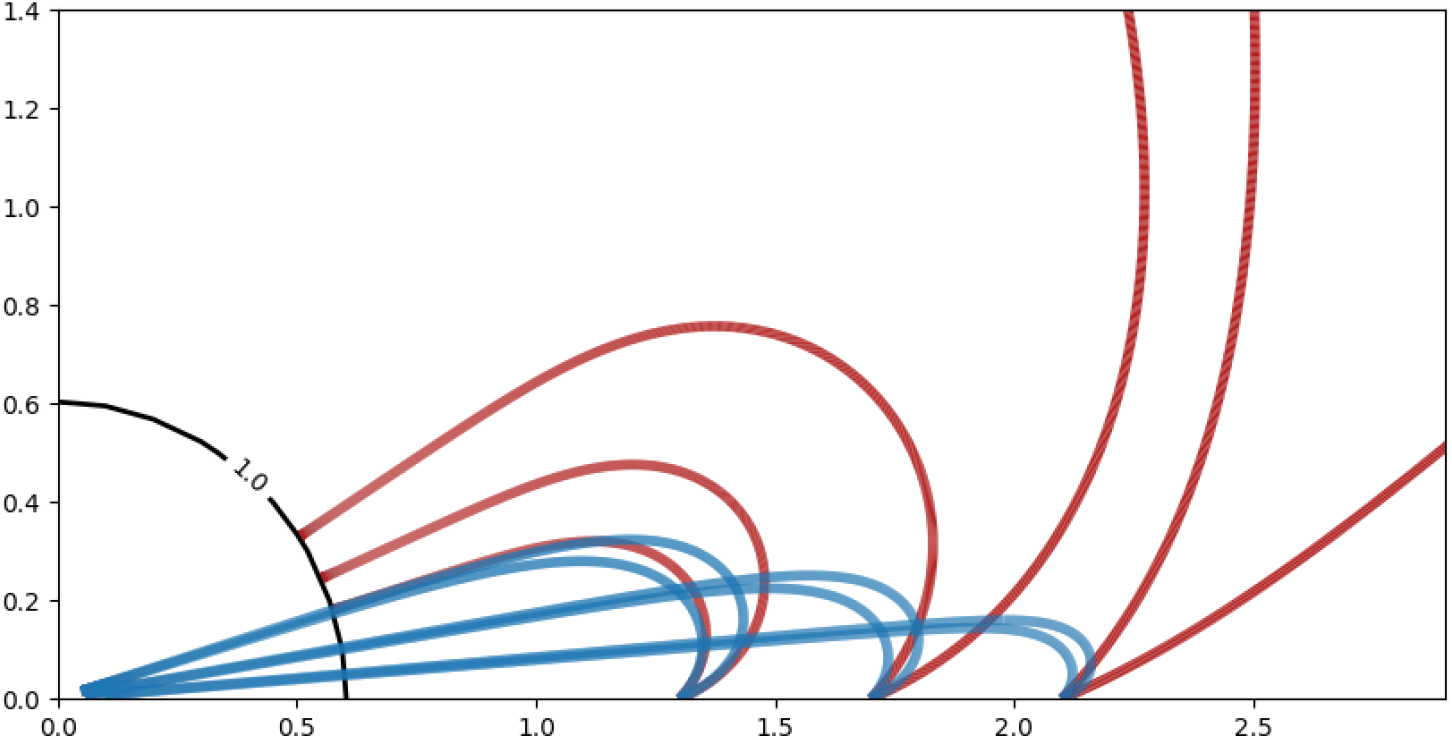
Trajectories of equation (1) (red) and equation (4) (blue) with a signal land-scape as in equation (2). The parameters of the response function (3) are given by *a*_1_ = *b* = 1, *c*_1_ = 2 and *d* = 5. Both sets of trajectories use the initial conditions *x*_0_ = 1.3, 1.7 and 2.1, *y*_0_ = 0, *θ*_0_ = *π/*4 and *θ*_0_ = *π/*8 (against the positive *x*-axis). The black circle indicates the level set of the signal strength *S*(*x, y*) = 1 where the trajectories of equation (1) terminate. Note that in these simulations we select the background velocity *v*_0_ = 0 in equation (4).

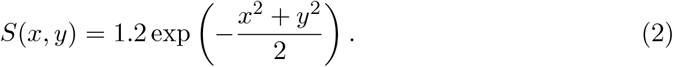

The response function *F* increases for vehicles of type 2 and decreases for vehicles of type 3. The sign “−” in the third equation of System (1) is chosen for an ipsilateral (type a) connection (the sensors influence the wheels on the same side), whereas the “+” sign is chosen for a contralateral (type b) connection (the sensors influence the wheels on the opposite side). The parameter *d* can be used to model the speed with which the agent changes its orientation in response to the signal gradient. In the following, we select the “−” in the third equation. The response function for a type 3a vehicle is chosen from the family

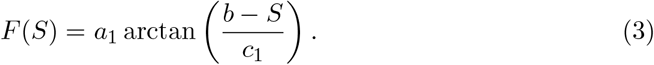

The parameter *a*_1_ *>* 0 determines the speed of the agent at low signal strength. The width of the velocity transition region is set by the parameter *c*_1_ *>* 0. In the setup of equations (1)-(3), it is necessary to assume as known a target level such that *F* (*S*^*^) = 0. This agent will also descend if it starts at a location with *S*(*x*_0_, *y*_0_) *> S*^*^.

We propose a different dependency of the velocity in the first two equations of (1) on ∇*S*. This is used to find critical points where ∇*S* = 0, although not all of those would necessarily be maxima of *S*. In this new model, the velocity is determined by a term *G*(||∇*S*(*x, y*)||^2^), where *G*(*s*) is a continuous increasing function with *G*(0) = 0. Thus, the gradient-search model becomes

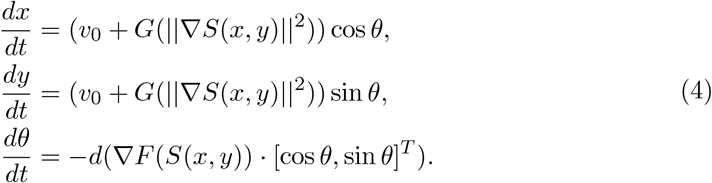

To avoid complete halt in the absence of a signal gradient, we also add a background velocity *v*_0_. In the following we use a second response function

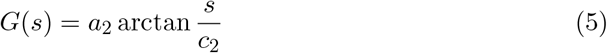

where we set *a*_2_ = 1 and *c*_2_ = 2 for our simulations. The numerical simulations of equations (1) and (4) in Figure 3 show that the dependence of the velocity on the signal gradient decreases the turning radius and thus allows the target to be found from a wider range of initial distances and directions.

## Results

Observations of *G. marleyi* individuals indicate a behavior that is not compatible with simple deterministic ODE models of Braitenberg vehicles. There are periods of motion and periods of rest [42]; see Figure 4, left panel. This leads us to introduce an energy reservoir *E*(*t*) ≥ 0 that is depleted during times of motion and is recharged during times of rest. During times of motion, *E* follows the differential equation

**Figure 4.**
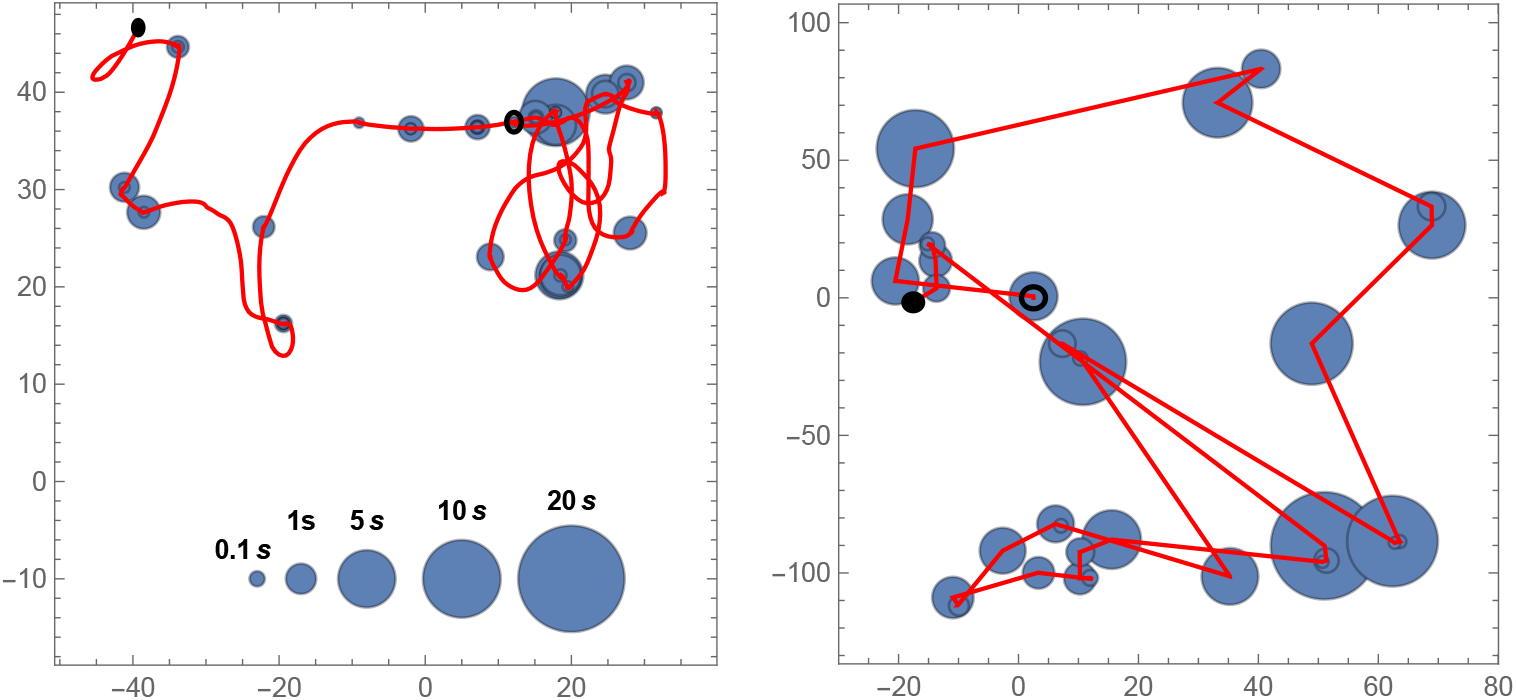
**(Left)** The recorded path of a *G. marleyi* for a duration of 40 seconds. Red lines show the movement while blue dots show locations at which the isopod rests. The size of the dots indicates the duration of the rest period. **(Right)** A sample path taken by the vehicle described by equation (4) in an empty environment, i.e. ∇*S*(*x, y*) = 0, and with Γ-distributed resting times. All coordinates are in millimeters. Starting and ending points of trajectories are marked with circles and disks, respectively.

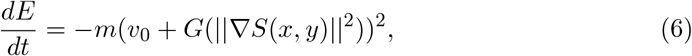

where *m* is a constant. Once *E* drops below a threshold *E*_*min*_, the movement stops and a resting time *T*_*R*_ is selected from a Γ-distribution. We advance the time to *t* + *T*_*R*_ and set the energy to *E*(*t* + *T*_*R*_) = *T*_*R*_, so that the recharging rate is 1, in appropriate units. Before the motion resumes, a new orientation *θ*(*t* + *T*_*R*_) is randomly selected from a normal distribution with standard deviation 1 centered at the last recorded orientation.

The empirical distributions of the times of rest and motion obtained from the video material are shown in Figure 5. The average rest and run times are 2.06 s and 3.81 s, respectively. To simulate resting times, we use a Γ-distribution with shape parameter *k* = 0.127 and scale parameter *θ* = 16.257. The average movement speed of the isopods while running is 3.04 mm s^−1^. We take that as the value of *v*_0_. A sample simulation of a vehicle exploring a flat environment is shown in Figure 4, right panel.

**Figure 5.**
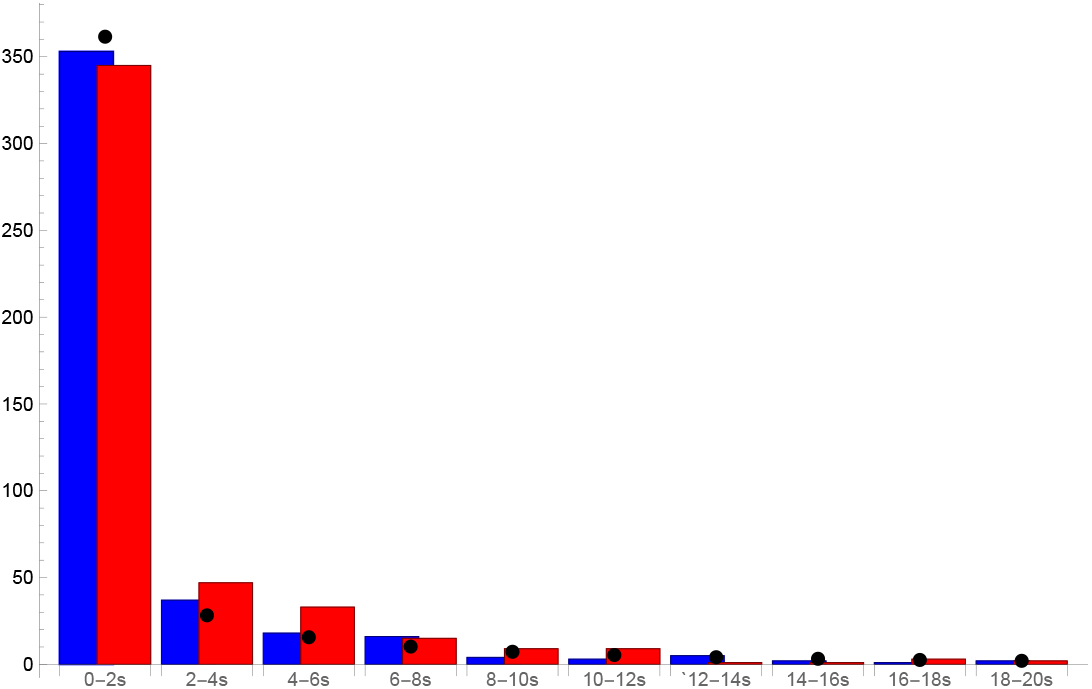
A histogram showing resting times (blue) and run times (red) of *G. marleyi* extracted from the video data. The black dots show the expected number of rest times in each interval from the Γ-distribution with parameters *k* = 0.127 and *θ* = 16.257. The isopod is considered to be resting if its velocity is below 0.05 mm s^−1^.

An example of this active searching behavior of a gnathiid in the olfactometer is shown in the left panel of Figure 6. In this trial, the gnathiid exhibits a behavior typically associated with a chemokenetic response to a chemical signal [43]. This gnathiid performs a series of directional changes, which is considered a host search strategy, while not successfully locating the source of the chemical signal. A sample simulation of an agent following the equation (4) in a landscape with multiple hills is shown in the right panel of Figure 6. Putting these two panels side by side should not be read as that we are trying to replicate the animals’ behavior in a very strict sense. Firstly, we are unable to reconstruct the chemical landscape in the olfactometer, despite the computational fluid mechanics simulations in [38]. Secondly, we do not have a wealth of observations, where we suspect a reaction to the chemical gradient, as seen by the smooth turns in Figure 6, left panel. Nevertheless, we see our simulations as valuable contributions toward a better understanding of the behavior of small aquatic ectoparasites in challenging chemical landscapes. Future research is needed to further solidify the parameter choices for equations (4)-(5).

**Figure 6.**
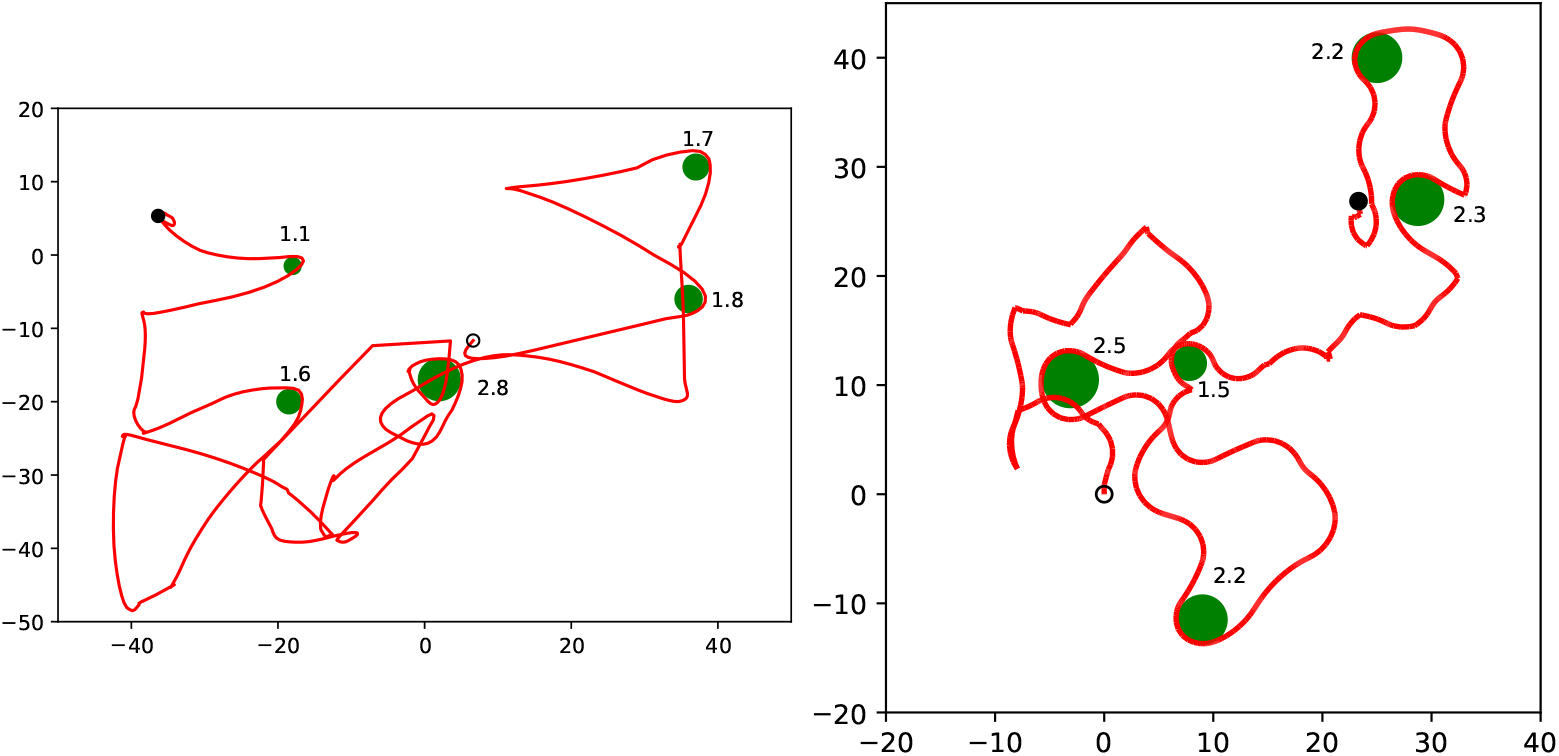
**(Left)** A recorded trajectory of a *G. marleyi* in the olfactometer. The source of the scent is roughly in the lower left corner. The starting point of the trajectory is marked by the circle, and the ending point is marked by the filled disk. The entire observation of the moving animal lasted about five minutes. Osculating circles at turning points are shown in green together with their radii in millimeters. The average radius is 1.8 mm. **(Right)** A simulation of an agent following equations (4)-(5) in a landscape of multiple peaks. The parameter values used for the simulation are as in the caption to Figure 3 and those following equation (5) except for *d* which is set to 0.5.

In contrast to *G. marleyi, O. faveolata* move continuously and do not stop. They also show much “rougher” trajectories than observed in the gnathiid specimens. *O. faveolata* exhibit a behavior associated with the chemotaxis response to a chemical signal [43], where the individual is attracted to the source of the signal. An example of this can be seen in Figure 7, where the individual successfully locates the source. Here we add a vigilance variable [44], 0 ≤ *r*(*t*) ≤ 1, which decays over time and modifies the equation that governs the direction *θ*. With these modifications, equation (4) becomes

**Figure 7.**
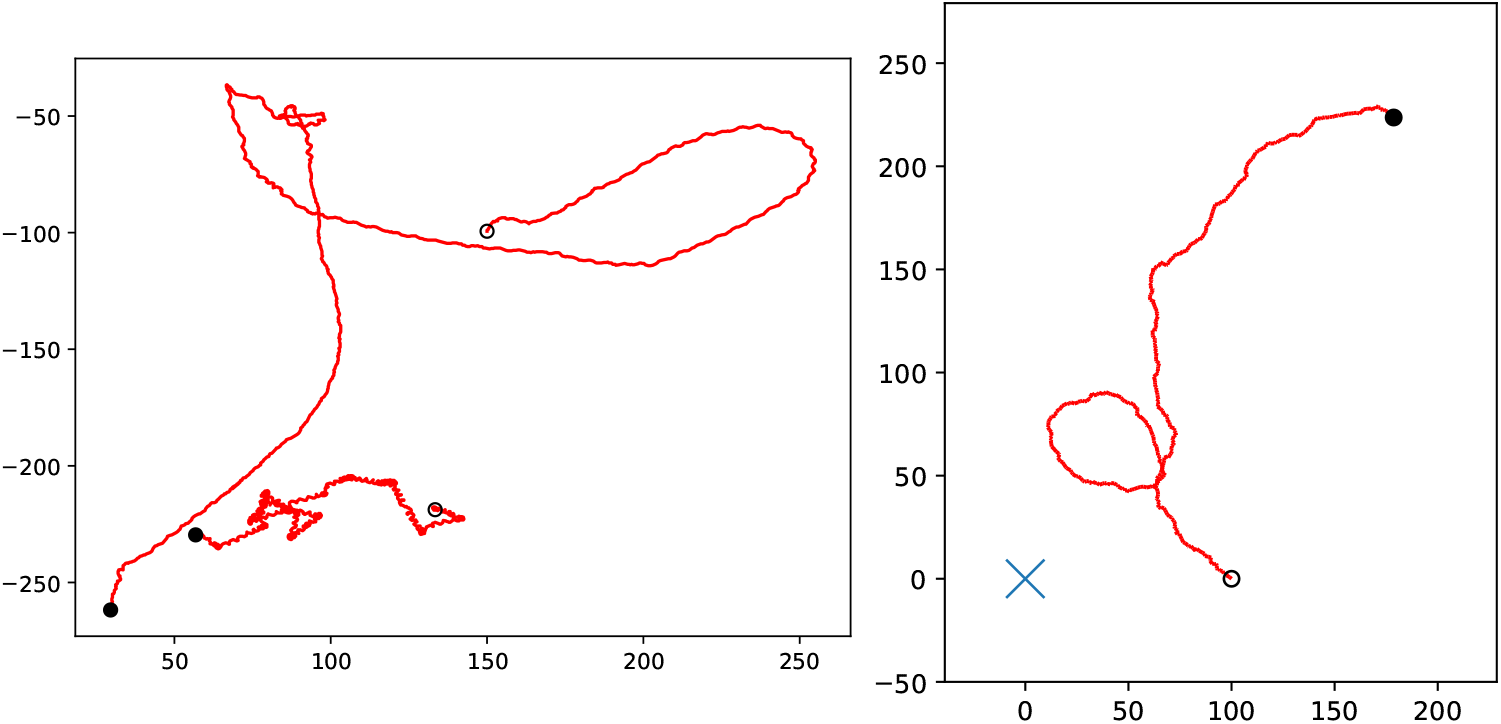
**(Left)** The recorded path of 2 coral larva specimens in the olfactometer. Positions are recorded at 5 Hz, and the coordinates are graded in millimeters. Starting and ending points are marked by circles and filled disks, respectively. Over the course of approximately three minutes, the animals decrease their distance from the source in the lower left corner. **(Right)** A simulated trajectory of a coral larva using equation (7) with parameters *a*_1_ = *a*_2_ = 100, *b* = 1, *c*_1_ = *c*_2_ = 2, *d* = 2, *σ*_1_ = *σ*_2_ = 5, *σ*_3_ = 1 and *v*_0_ = 0.15. The peak of the signal *S* is marked with a blue cross.

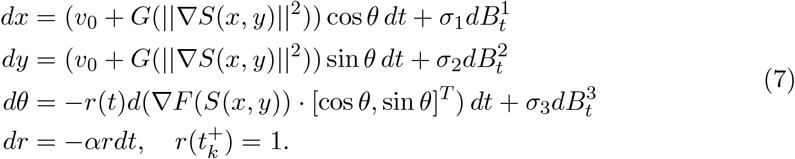

At time points *t*_*k*_ chosen from an exponential distribution with parameter *β* = 1, we set 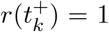, thus renewing the vigilance to its maximal value. Here, *α* = 0.5 is the rate at which the interest is lost. *B* denotes a ℝ^3^-valued Wiener process [45]. This system is solved numerically using the Euler-Maruyama method [46]. The result is shown in the right panel of Figure 7.

## Discussion

Biologists, cognitive neuroscientists and engineers are conducting exciting research on the search behavior of animals and the possible replication of those behaviors in *in silico* agents or in robots [10, 11, 15–17, 47, 48]. [11] have complemented observations of larval and adult fruit flies *Drosophila melanogaster* with Braitenberg vehicles of types 3a and 2b, respectively. To the best of our knowledge, planktonic animals have not yet served as a biological source of inspiration. Here we present observations of both gnathiid isopods and coral larvae and pair them with simulations of type 3a Braitenberg vehicles following novel mathematical models. A rich field of contemporary applications of Braitenberg vehicles is robotics, where agents are programmed to perform a specified tasks. Observations of freely moving animals in the custom-made olfactometer, as done here, show behaviors that require modifications of the deterministic models, such as those in equations (1) and (4). New in this work is the use of the signal gradient ∇*S* in equation (4) as the driving force of velocity. In nature, signals exist on different background strengths, for example, the overall density of the host fish population. Moreover, beyond the stochastic versions of previously deterministic ODEs that arise from noisy sensors [49, 50], our observations of *G. marleyi* show frequent changes between motion and rest. This leads us to introduce an energy reservoir following the equation (6). Even when an animal is suspected of following the chemical signal, there is a distinct possibility that an individual is losing vigilance [44] after a certain period of time, see

Figure (6), left panel, and equations (7). This could be due to neurosensory limitations, physical fatigue, and the need to rest between bouts.

Gnathiid isopods adopt similar host-finding strategies as other blood-feeding arthropods. *G. marleyi* likely uses chemical signals to position itself in locations near hosts or detect approaching hosts. Then, as a host nears them, visual or hydrodynamic cues become important. This can be seen in their behavior within the olfactometer, with chemical signals simulating a searching behavior indicated by their overall higher levels of activity than when in water with no chemical signals. From a mathematical point of view, the difference between the presence and absence of a chemical signal is similar to the difference between a biased and an unbiased random walk, respectively; see Figure 8. However, in the absence of other stimuli that indicate a nearby host, *G. marleyi* show long periods of rest.

**Figure 8.**
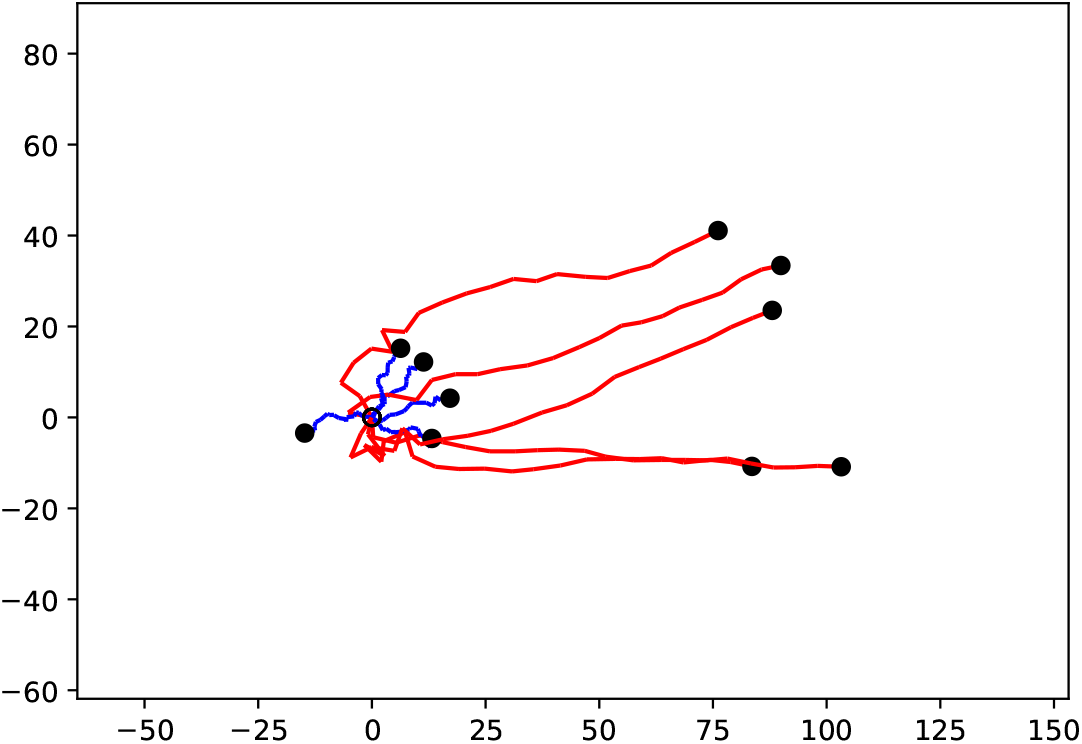
Two sets of trajectories of equation system (7) with *S*(*x, y*) = *x* (red) and *S*(*x, y*) = 0 (blue). All trajectories start at the origin with randomized initial directions (open circle) and the end of each trajectory is marked with a black dot. The agents with a signal quickly turn to the right, while the agents in an empty environment only perform a localized random walk.

Many animals respond to a multitude of sensory stimuli with different intensities. For example, an animal may respond weakly to a broad scent or a chemical signal but strongly respond to a more focused visual or hydromechanical signal. To model this, we modify the third equation from the system (7) and include the presence of a second signal *R* = *R*(*x, y*). This results in

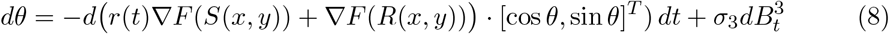

The second signal *R* is chosen to be much narrower in dispersion than *S*. However, the agent’s response to the second signal is not affected by decreasing vigilance *r*(*t*). For gnathiids, this can be interpreted as a visual or hydromechanical signal in the immediate vicinity of the fish host. Figure 9 shows that this modification prevents the agent from leaving the region surrounding the signal peak, where *R*(*x, y*) *> S*(*x, y*) as the ∇*F* (*R*(*x, y*)) term causes it to turn much more rapidly. However, it does not significantly improve the agent’s ability to reach this region. Naturally, an animal seeking a host or a mate would eventually dock onto its target.

**Figure 9.**
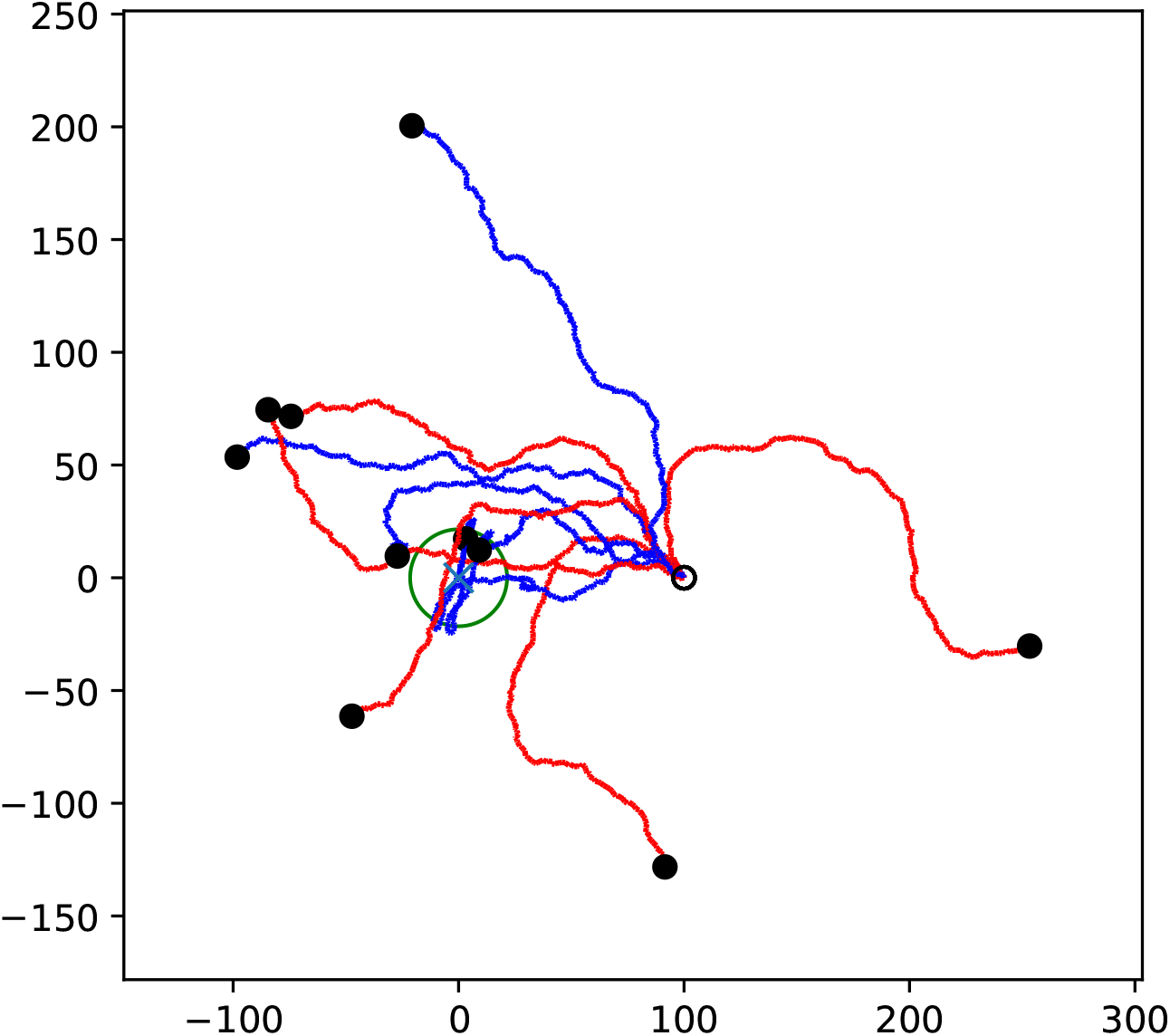
Ten simulations using the vigilance models (7) (red) and (8) (blue). The initial value for all simulations is (*x, y, θ, r*) = (100, 0, 3*π/*4, 1), with the position indicated by the black circle. The times of renewal *t*_*k*_ are selected from an exponential distribution with parameter *β* = 1. The green circle indicates where the second signal exceeds the first, that is, *R* ≥ *S*.

Chemical signaling landscapes in nature are rarely static. Either the source moves, or the carrying fluid moves, such as wind [48] or the ocean water in a coral reef. To implement a source moving rightward at a constant velocity, we change the signal landscape in equation (2) by replacing *x* by *x* −(*tv*_1_ − *x*_0_), where *v*_1_ is the speed at which the landscape moves and *x*_0_ is the initial location of the center or peak. In the simulations, the agent starts at *y*_0_ = 0 below the path of the source on the line *y* = 20 mm; see Figure 10. The center of the landscape begins above and to the left of the agent. The simulation ends if the agent comes within 3 mm of the target. As shown in the examples in Figure 10, when *v*_1_ ≤ *v*_0_ (that is, the speed of the agent when it has no signal), the agent is able to frequently successfully reach the center of the landscape. The searches fail when *v*_1_ *> v*_0_ as the agent falls behind. Most isopods are known to hunt at night or at crepuscular times. In addition to a reduced risk of predation by planktivorous fish, hosts are then easier to target because they are quiescent. Moreover, in shallow water (*<* 5 m), there is usually less wind effect at night, and thus the water is calmer.

**Figure 10.**
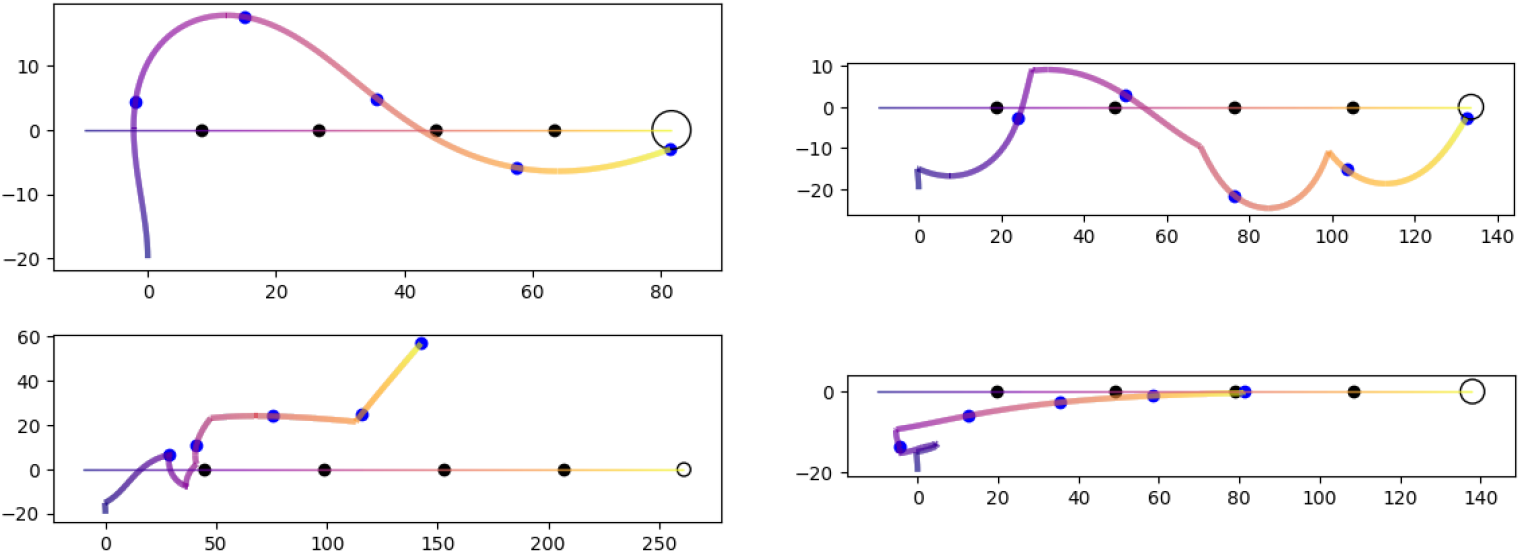
Four simulations showing an agent pursuing a source moving from left to right indicated by the horizontal line. The colors of both paths indicates equal time stamps and dots mark the passage of every fifth of the total time. A simulation ends successfully when the vehicle enters the circle of 3 mm radius at the terminal point. The top two panels show successful pursuits where the source moves slightly slower than the agent’s passive speed *v*_0_. The second pair show failed pursuits where the source moves slightly faster than the agent’s passive speed.

In addition to the use of Braitenberg vehicles as a tool for studying the behavioral responses of tiny metazoans to chemical signals, we have also gained valuable information to further refine the development of the olfactometer. Originally invented to study the olfactory responses of small insects [51], the olfactometer has also been effectively used to record the behavior of aquatic organisms. However, slower or constantly moving organisms may be more ideal candidates for future studies. Gnathiid isopods, as sit-and-wait predators, appear to rely on other sensory cues during their host finding, which are intentionally not present in the olfactometer. This absence may affect their true host-finding behavior. With respect to the construction of the olfactometer, a key limitation is that the chamber used to introduce the organism renders a large portion of the arena invisible. If possible, this should be modified to remove this obstruction or reduce it in size as much as possible. Other minor structural recommendations include using a material that does not rust as easily to prevent potential cue contamination, digital flow meters to better assure accurate and even flow rates, and a redesign to reduce weight to make travel with the olfactometer easier.

## Acknowledgments

PH is grateful for the kind hospitality of the Center for Information Services and High Performance Computing (ZIH) at the Dresden University of Technology in Dresden, Germany, during a sabbatical. Special thanks are given to Professors Wolfgang Nagel and Andreas Deutsch. The authors thank Amber Packard of the Environmental Analysis Laboratory at the University of the Virgin Islands and Andrew Baker, Katherine Hardy, and Maren Stickley at the Rosenstiel School of Marine, Atmospheric and Earth

Sciences at the University of Miami for use of facilities and assistance with laboratory and field work. The authors also thank J. Rudi Strickler, Kyle Jansson, and the University of Wisconsin - Milwaukee Prototyping Center for their development of the aquatic olfactometer and assistance with troubleshooting. We thank two unknown readers for valuable comments on the manuscript.

